# Linking lineage and population observables in biological branching processes

**DOI:** 10.1101/527291

**Authors:** Reinaldo García-García, Arthur Genthon, David Lacoste

**Affiliations:** Gulliver, ESPCI Paris, PSL University, CNRS, 75005 Paris, France

## Abstract

Using a population dynamics inspired by an ensemble of growing cells, a set of fluctuation theorems linking observables measured at the lineage and population levels are derived. One of these relations implies inequalities comparing the population doubling time with the mean generation time at the lineage or population levels. We argue that testing these inequalities provides useful insights into the underlying mechanism controlling the division rate in such branching processes.

## I. INTRODUCTION

The question of how a cell controls its size is a very old one [1], which despite decades of research is still under intense focus, because the old experiments have only provided incomplete answers while a new generation of experiments based on the observation and manipulation of single cells in microfluidic devices is becoming more and more mature [2]. For instance, with time-lapse single cell video-microscopy, entire lineages of single cells such as *E. coli* can be traced over many generations. These experiments allow to investigate mechanisms of cell size control (cell size homeostasis) with unprecedented statistics both at the single cell level and at the level of a population.

Many policies of cell size control have been introduced: the “sizer” in which the cell divides when it reaches a certain size, the “timer” in which the cells grows for a specific amount of time before division, and the “adder” in which cells add a constant volume each generation [3]. The adder principle is now favored by many experiments [4–7], yet there is no consensus on why a specific regulation emerges under certain conditions, and how it is implemented at the molecular level.

Another important question is how to relate measurements made at the lineage and at the population levels. A classical study revealed the discrepancy between the mean generation time and the population doubling time [8] in an age-dependent branching process with no mother-daughter correlations, called Bellmann-Harris process in the literature on branching processes [9]. Importantly, it is still not known at present how to relate the mean generation time and the population doubling time in general models of cell size control.

Inspired by single-cell experiments with colonies of prokaryotic cells in microfluidic devices [5, 10], we consider here continuous rate models (CRM), based on stochastic differential equations [4, 11]. Unlike discrete stochastic maps (DSM) [3], there is no need in CRM to rely on a policy function since the division mechanism is encoded in the functional form of the division rate. The corresponding population dynamics has an interesting thermodynamic structure uncovered in Refs. [12, 13], which we also exploit here to derive three new fluctuation relations. As usual with fluctuation theorems [14], our results map typical behaviors in one ensemble (here the population level) to atypical behaviors in another one (here the single lineage level). A similar connection lies at the basis of an algorithm to measure large deviation functions using a population dynamics [15, 16]. In the mathematical literature on branching processes, relations of this kind are known as Many-to-One formulas [17]; they explain the existence of a statistical biais, when choosing uniformly one individual in a population as opposed to following a lineage.

This paper is organized as follows: In the next section, we introduce two CRM dynamics, which will be studied in this paper, namely a size-controlled and an age-controlled model. In Sec. III, we derive a fluctuation relation for the first type of models. This fluctuation relation maps the single lineage level and the population level. We test the relation numerically, and we derive related inequalities between the mean generation time and the population doubling time. In the next section, Sec. IV, we derive a second more general fluctuation relation, valid for both size models and age models without correlations. We also explain how this framework is related to the notion of “fitness landscape” introduced in Ref. [18]. Then, we analyze age models with correlations between mother and daughter cells. Finally, we conclude in Sec. V, while all the important technical details are given in the appendices.

## II. CONTINUOUS RATE MODELS

Let us consider a population of cells as shown in Fig. 1a, which grow by division into only two offsprings at the end of each cell cycle. This population dynamics can be studied at three distinct levels: the lineage level (red), the population snap-shot (blue) and the tree level which includes the complete phylogeny [19]. For bacteria such as E. coli growing in a rich medium, each cell cycle is well described by an exponential growth phase [20], which for the cell cycle *i* can be parametrized byonly three random variables shown in Fig. 1b: the size at birth 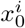, the growth rate *ν*^*i*^ and the generation time *τ*_*i*_.

**FIG. 1.**
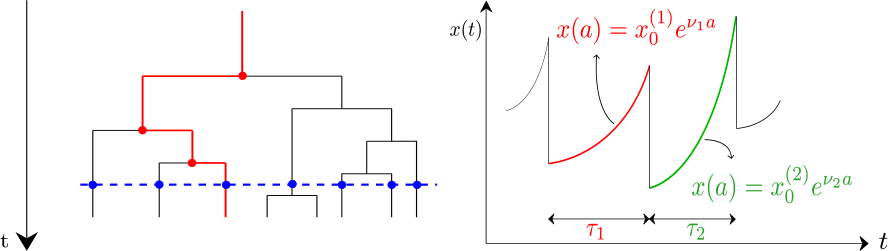
(a) Representation of the three main levels of description of the ensemble of cells: the lineage level (red), the population snapshot (blue) and the entire tree (black). (b) Evolution of the cell size *x*(*t*) along a lineage. The cell cycle *i* is parametrized by the generation time *τi*, the growth rate *νi* and the size at birth 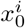.

In the following, we consider successively two types of continuous rate models: in the first one based on size, the division rate depends only the size and in the second one, the division rate depends only on the cell age.

### A. Size-dependent division rate

Let us first consider a model with size-dependent division rate. The evolution of the number of cells of size *x* and single cell growth rate *ν* at time *t*, *n*(**y**, *t*) with **y** = (*x*, *ν*), obeys the equation [4, 11]:

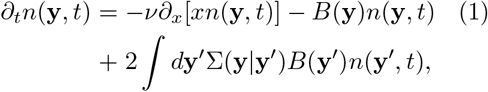

where *B*(**y**) is the division rate and Σ(**y**|**y′**) is the probability for a newborn cell to have parameters **y** given that the mother cell has parameters **y′**. By integrating Eq. (1) over **y**using the condition ∫ *d***y Σ**(**y|y′**) = 1, a deterministic equation of evolution of the total population *N* (*t*) = ∫ *d***y***n*(**y**, *t*) is obtained.

The instantaneous growth rate of the population is defined as 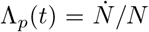, while the growth rate of the total volume of the cells is 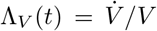 with *V* (*t*) = ∫ *d***y***xn*(**y**, *t*). When a steady state for the variable **y** is reached, both Λ_*p*_ and Λ_*V*_ become independent of time and equal to each other [19].

If instead of the full population, we consider the dynamics at the lineage level, the natural quantity to study is the *probability density* of the cell to have size *x* and growth rate *ν* at time *t*, *p*(*x*, *ν*, *t*), which satisfies the evolution equation

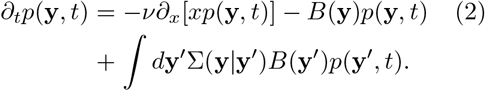

Note the difference with Eq. (1) due to the absence of the factor 2 in front of the integral, rendering *p*(**y**, *t*) normalizable at any time, ∫ *p*(**y**, *t*)*d***y** = 1.

### B. Age-dependent division rate

When the division rate depends on the age of the cells instead of their size, the structure of the model is rather different from that of the previous subsection. Let us now introduce a further distinction between two types of age models. In the first type, the interdivision times of mother and daughter cells are uncorrelated, and the division rate is determined by the age of the cells only. Such a model is usually termed independent generation times (IGT) model or Bellmann-Harris process [9]. In a second type of models, the division rate may depend on other variables besides the age, and as a result, mother-daughter correlations will be present.

In the case of the IGT type of models, the density of cells having age *a* in the population at time *t*, *n*(*a*, *t*), satisfies the evolution equation

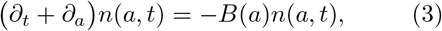

with the boundary condition:

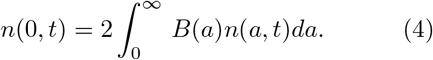

As before, *B*(*a*) denotes the age-dependent division rate. The physical interpretation of the boundary condition (4) is clear: Each dividing cell gives rise to two newborn cells (i.e. two cells with age *a* = 0). The total number of cells in the population at time *t* follows by integration of the density, *N* (*t*) = ∫ *n*(*a, t*)*da*.

As in the case of size control, lineage dynamics can be directly encoded in the evolution of the age distribution. Such dynamics reads

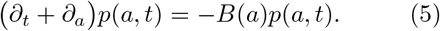

which is complemented by the boundary condition:

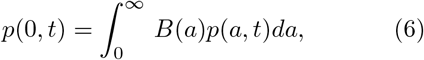

so that probability is conserved and *p*(*a*, *t*) is normalized.

In a second type of models, correlations in the inter-division times are accounted for by adding an extra dependence of the division rate on the growth rate, *B*(*a*, *ν*), while introducing at the same time correlations between the growth rate of mother and daughter cells. The model then reads

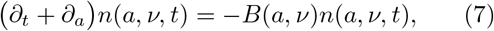

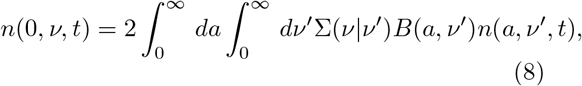

at the population level, and

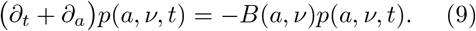

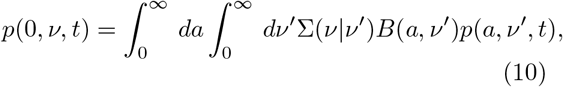

at the lineage level.

## III. FLUCTUATION THEOREM FOR DYNAMICAL ACTIVITY

We now address the problem of connecting lineage to tree or population snapshot statistics in models with size control. The evolution of a given cell from time 0 to the time *t* is encoded in the trajectory 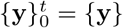.

For the case of size-controlled model, we derive in Appendices A, path probabilities representations at the population and lineage levels, which are given by (A10) and (A9) respectively. Comparing these two expressions, we see that a possible way to bring both distributions “closer” together, is to multiply the division rate at the lineage level by the factor *m*, and to consider a lineage starting from the same initial condition as that of the population.

Then, we introduce the dynamical activity 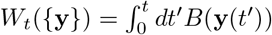, which quantifies the activity of cell divisions, and the time averaged population growth rate

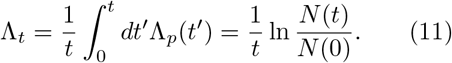

After multiplying the relation mentioned above between path probabilities by an arbitrary trajectory-dependent observable *A*(*x*, *ν*), and after taking the average, for the special case where *m* = 2, one obtains the following fluctuation relation:

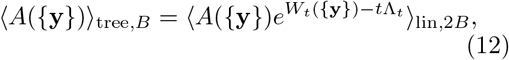

where 〈‥〉 _tree,*B*_ denotes a tree average generated by the original dynamics with a division rate *B* while 〈‥〉 _lin,2*B*_ denotes a lineage average with a modified dynamics that has a division rate 2*B*. The reason for this modified division rate is that each cell divides into *m* = 2 cells, as a result a factor two appears at the population level in Eq. (1), which is absent for the corresponding equation at the lineage level. In the particular case where the observable *A* only depends on **y**(*t*) instead of the full-trajectory {**y**}, Eq. (12) relates the lineage level to the population snapshot level instead of the tree level. The mapping also requires that the original and the modified dynamics start with the same initial condition **y**(0), defined here in terms of cell size and growth rate.

For the specific choice *A*({**y**}) = *δ*(*W* − *W*_*t*_({**y**}), Eq. (12) leads to Crooks-like relation [14]:

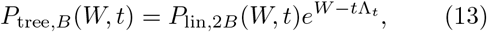

which relates the distribution of dynamical activity at time *t* in a tree (resp. in a lineage): *P*_tree,*B*_ (*W*, *t*) (resp. *P*_lin,2*B*_ (*W*, *t*)). This relation is illustrated in Fig. 2 for a population of cells growing with a constant single cell growth rate *ν*. Numerically, instead of working directly with Eq. (1), we simulate an equivalent Langevin equation, which accounts for deterministic growth with the rate *ν* and stochastic cell divisions with a rate *B*(*x*, *ν*). In the simulation, the division has been assumed to be symmetric and the single cell growth *ν* constant, which corresponds to the particular choice of Σ(**y|y′**) = *δ*(*ν* − *ν′*)*δ*(*x* − *x′*/2). Note that this dynamics bears some similarity to that of stochastic resetting introduced in Ref. [21], with the difference that in our case the resetting of the size is relative to the current size before division, while in this reference the resetting was to a constant position. Another important difference is the absence of diffusion in our model.

We have used normalized units of time and size, so that *ν* = 2 and *B*(*x*, *ν*) = *νx* in these units. Since Λ_*p*_ = Λ_*V*_ = *ν*, Λ_*t*_ = 2, the two distributions. measured at the time *t* = 2 cross as expected at *W* = 4 (top figure). The bottom figure confirms that the slope of the log-ratio of the two probability distributions is indeed −1 as expected from Eq. (13).

Let us emphasize the following points concerning our first main result: This fluctuation relation is very general, it holds whether or not the single cell growth rate fluctuates, i.e., for arbitrary forms of the kernel Σ and arbitrary division rate *B*(*x*, *ν*). There is no requirement that the population should be stationary neither at time 0 nor at time *t*. Further, it generalizes to the case that each cell has *m* offsprings instead of two, provided that this number *m* is independent on the state of the system **y** and that the modified lineage dynamics has a division rate *mB*(*x*, *ν*), as shown in Appendix A 3.

The normalization of *P*_tree,*B*_ (*W*, *t*) in (13) leads to the relation:

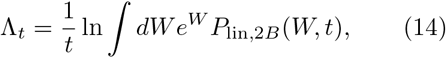

which could be used either to infer the population growth rate from lineage trajectories or to infer the form of the division rate *B* using lineage and population trajectories [22]. In the next subsection below, we provide such a numerical illustration.

**FIG. 2.**
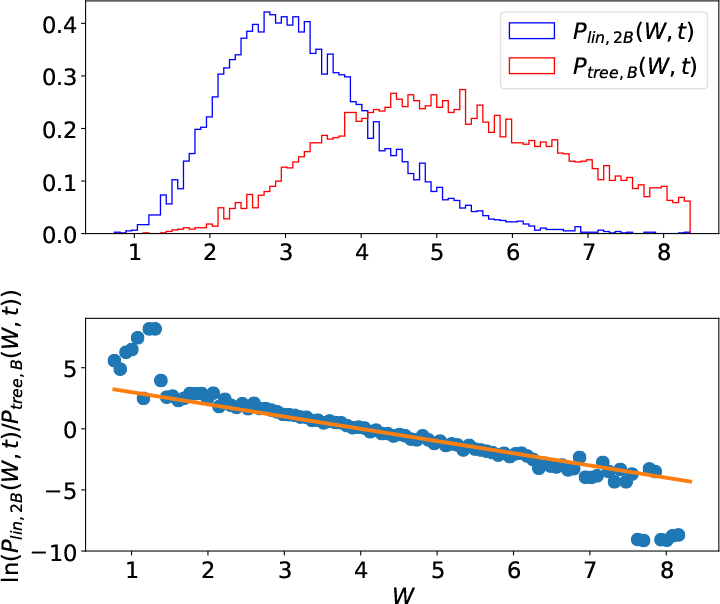
Illustration of the fluctuation relation in the case of growth with a constant growth rate *ν* = 2, showing the distributions of dynamical activity at the time *t* = 2 in a tree and in a lineage (top figure) and the log-ratio of these probability distributions (bottom figure).

### A. Application to the determination of a population growth rate

Since the variability of single cell growth rate is known to be important experimentally [20], we now discuss its role on the population growth rate in light of our results. A simple way to study this question in a simulation is to assume that the single cell growth rate *ν* is distributed according to a normal distribution of mean *ν*_*m*_ and variance *σ*_*ν*_. Since *ν*_*m*_ and *σ*_*ν*_ take the same value at each division, there is no correlation between the generation time of the mother and daughter cell. This is the situation studied in Fig. 3, where the population growth rate Λ_*p*_ is plotted as function of *ν*_*m*_. In the absence of variability where *σ*_*ν*_ = 0, we have Λ_*p*_ = *ν*_*m*_, which is shown as a black dashed line in the figure. In the presence of variability, this figure confirms that the growth rate of the total volume Λ_*V*_ equals the growth rate of the population where both of them have been measured from the statistics of the final population at a fixed time. Importantly, such a determination of the population growth rate also agrees (within errors bars) with the one based on the fluctuation relation of Eq. (14) using lineage trajectories. Therefore, this shows that the fluctuation relations of Eq. (14) could be used as a numerical method to determine a population growth rate based on lineage statistics.

Another striking feature of Fig. 3 is that regard-less of the determination of Λ_*p*_, all the points are below the dashed line. The interpretation is that in a snapshot at time *t*, it is less likely to see cells with a short generation time (corresponding to large single cell growth rates), therefore the distribution is biased towards small single cell growth rate [19]. Since the population growth rate generally increases with respect to the single cell growth rate *ν*_*m*_, this bias leads to a decrease of the population growth rate with respect to the case of no variability in the single cell growth rate.

**FIG. 3.**
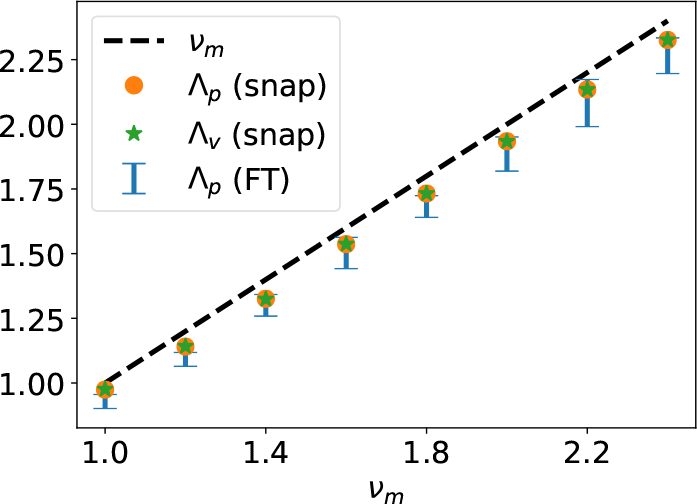
(a) Population growth rate Λ_*P*_ versus the mean single cell growth rate *ν*_*m*_: from a population snapshot (orange circles), from the growth rate of the total volume (green stars), and from the fluctuation relation of Eq. (5) of the main text. Here the cell growth rate is taken from the normal distribution 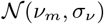 and *B*(*x*, *ν*) = *νx*. Error bars have been obtained by using the fluctuation relation on 1000 trajectories and then repeating the estimation another 50 times.

As mentioned in the introduction, the fluctuation relation of Eq. (12) includes in itself a statistical biais: when choosing uniformly one individual in a population, an individual belonging to a lineage with prolific ancestors is more likely to be chosen, as a result, the jump rate on a lineage must be multiplied by the mean number of offsprings. Although variability in the single cell growth rate also introduces a form of statistical bias as explained above, the biais is not exactly the same one as that contained in the fluctuation relation. In any case, we would like to point out a comprehensive theoretical study on the effect of variability on the population growth rate, namely [23]. This study confirms that in the case of size models with i.i.d. single cell growth rates, variability indeed lowers the Malthusian growth rate as observed in figure 3. This work also discusses age models, with and without correlations in single cell growth rates, and concludes that in general, variability may lead to either a positive or negative trend on the population growth rate.

**FIG. 4.**
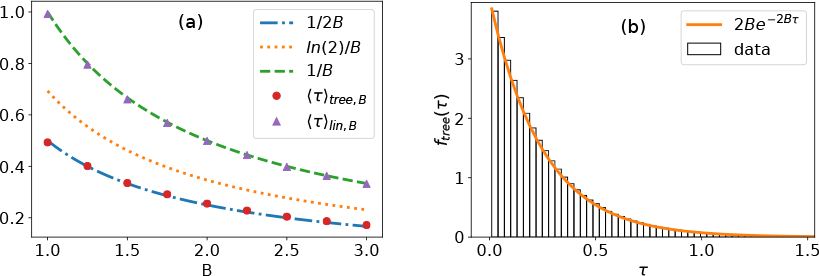
(a) Mean generation times evaluated in a tree (red circles) and in lineage (violet triangles) against the division rate *B*. Theoretical predictions are shown as dashed lines and the doubling time *T*_*d*_ = ln(2)*/B* is shown as a dotted line. (b) Distribution of generation times in the tree *f*_tree_(*τ*). In these figures, the division rate, *B* and the single cell growth rate, *ν*, are constant and equal to each other.

### B. Consequences for the distribution of generation times

An important quantity in population dynamics is the distribution of generation times *f* (*τ*). This quantity can be evaluated from the observable [24]:

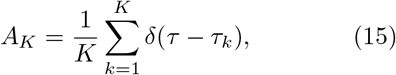

where the index *k* runs over all the *K* cell cycles which have appeared in the trajectory that starts from *t* = 0 to final time *t*. This observable can be evaluated either on a lineage or on a tree. By reporting *A*_*K*_ as the observable *A* in Eq. (12), one deduces the relation

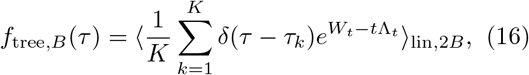

where a summation over the random variable *K* and a dependence on the final time *t* are implicit. In the particular case where the division rate *B* is constant, *W*_*t*_ = *t*Λ_*t*_ and therefore *f*_tree,*B*_ (*τ*) = *f*_lin,2*B*_ (*τ*). In this case, the generation time distribution in a lineage is the simple exponential *f*_lin,*B*_ (*τ*) = *B* · exp (*Bτ*) with mean 1/*B*. It follows that *f*_tree*,B*_ (*τ*) = 2*B* · exp (2*Bτ*) with mean 1/(2*B*). Fig. 4 confirms that the distribution of generation times has the expected properties.

## IV. A SECOND FLUCTUATION THEOREM TO RELATE LINEAGE AND TREE STATISTICS

### A. Size-controlled model

When the division rate *B* is not constant, the distribution of generation times will no longer be exponential, but we may still wonder how mean generation times observed at the lineage and tree levels compare to each other. In order to address this issue, we derive a different fluctuation theorem that connects this time the lineage and tree statistics with the *same* division rate *B*. More precisely, it follows from a direct comparison of (A10) and (A9) taking again *P*_0_ = *p*_0_. Since the division rate is the same in both probability distributions, we stick to the notations introduced above, except that now the index *B* will be omitted.

This allows to write

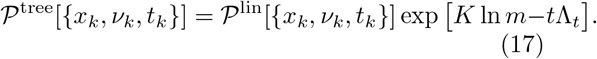

Now, by multiplying the above relation by an arbitrary trajectory-observable *A* and taking *m* = 2, we obtain:

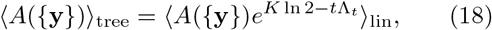

where *K* = *K*({**y**}) counts as in Eq. (15) the number of divisions.

In the particular case where *A*({**y**}) = *δ*(**y** − **y**(*t*))*δ*_*K*,*K*(*t*)_, Eq. (18) leads upon averaging, to a relation between the joint probability distributions of size, growth rate and number of divisions at the lineage and tree levels [18]:

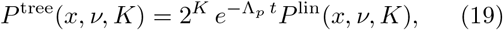

which we call a *local* fluctuation relation. Averages over lineages within a population can be carried out with respect to either a chronological or to a retrospective distribution [12, 13, 24], which correspond respectively to our lineage and tree probability distributions. Let us briefly comment on a connection to a discussion presented in Ref. [18]. Elimination of *K* in Eq. (19) leads to a fluctuation theorem only involving phenotypic traits *x* and *ν*:

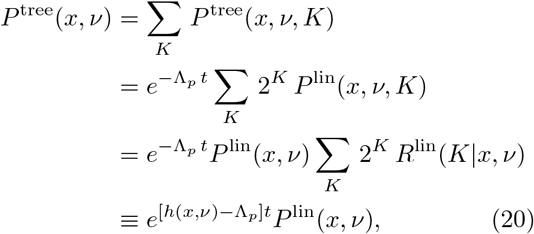

where we have introduced the probability of the number of division events conditioned on size and growth rate at the lineage level, *R*^lin^(*K*|*x*, *ν*) and the equivalent of the “fitness landscape” of Ref. [18] reads

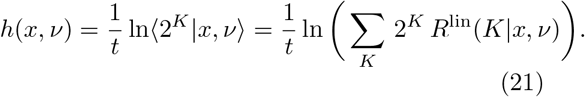

By summing over *K* in Eq. (19), one obtains

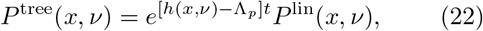

in terms of a function *h*(*x*, *ν*) called “fitness landscape” in Ref. [18]. Eqs. (19)–(22) show that the knowledge of the two phenotypic probability distributions *P*^tree^ and *P*^lin^ can be used to infer a fitness function for size and growth rate.

### B. Consequences for the generation times

Let us also introduce the Kullback-Leibler divergence between two probabilities *p* and *q*:

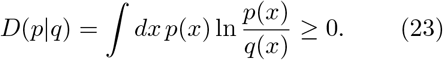

Using the fluctuation relation of Eq. (17), we obtain

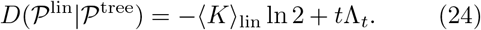

On large times *t*, we can use the relation ⟨*τ*⟩_lin_ = *t*/⟨*K*⟩_lin_, which together with the definition of the population doubling time *T*_*d*_ = ln 2/Λ_*t*_, leads to the right inequality in

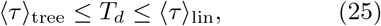

while the left inequality follows very similarly using 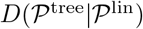.

In the case that *B* is constant shown in Fig. 4a, Eq. (25) is trivially satisfied. For *B* non-constant of the form *νx*^*α*^, the inequalities are verified numerically in Fig. 5. This figure shows that the mean generation time for lineage (resp. tree) approaches the doubling time in the limit of large *α*, because in this limit the distribution of generation times becomes peaked at *T*_*d*_.

**FIG. 5.**
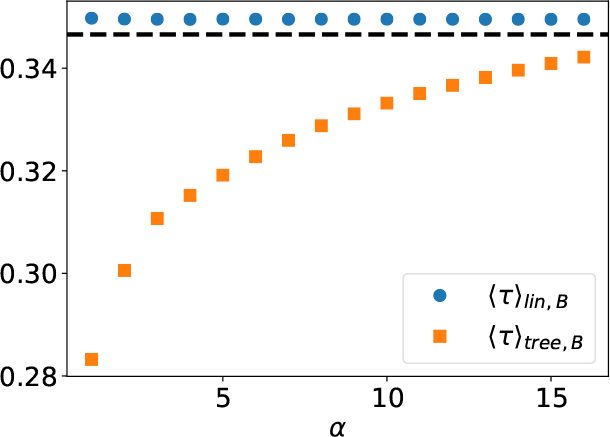
Mean generation times measured in a lineage or in a tree versus the exponent *α* entering in the division rate *B*(*x*, *ν*) = *νx*^*α*^. The dashed line represents the doubling time ln 2/Λ, the single cell growth rate is *ν* = 2, and values *α* ∈ [1, 16] are shown.

### C. Age-controlled model

Beyond cell size control models, one can also consider age models, which may have or not mother-daughter correlations. Let us first consider the case where correlations are absent, the so-called IGT model, and let us focus on the distribution of generation times either in a lineage or in a population.

As in the case of size control, lineage dynamics of age-structured models can be directly encoded in the evolution of the age distribution, as prescribed by Eqs. (5) and (6). Let us consider steady-state conditions:

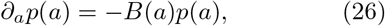

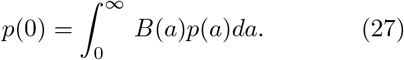

A nice feature of age models is that the generationtime distribution can be accessed directly. This is so because generation time distribution is the age distribution of the dividing cells. We proceed to compute this distribution for individual lineages in age-structured IGT models. First, note that from (26) immediately follows that

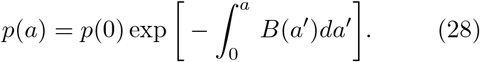

Relying on the relation between generation time distribution and age distribution of *dividing* cells, we can write

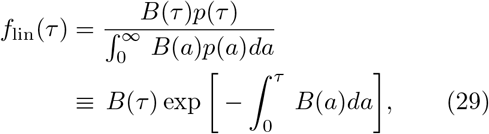

where we have used (27) and (28).

Now in order to obtain the distribution of generation times at the population level, we start from Eqs. (3) and (4). Again, we focus on stationary conditions for which the total number of cells in the population grows exponentially, as 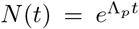. In that case, de density can be written in terms of the stationary probability density of cells with a given age as 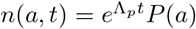, where *P*(*a*) is the stationary age distribution of the population. We have:

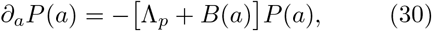

with the boundary condition:

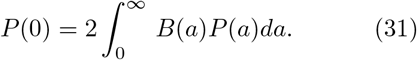

It is worth noting that normalization of *P*(*a*) in (30), leads, using (31), to the following identity

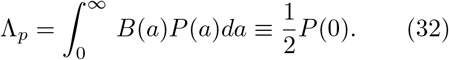

We can now proceed to compute the generation time distribution, by computing the age distribution of dividing cells. We have first for the stationary distribution from (30):

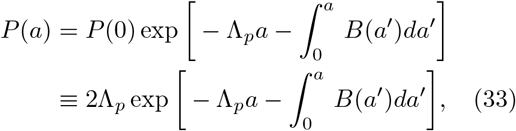

where we have also used (32). On passing by, we highlight an important relation for IGT models obtained from the normalization of *P*(*a*) in Eq. (33):

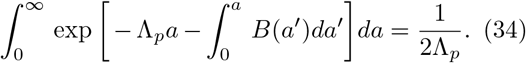

We can now calculate the generation time distribution, which reads

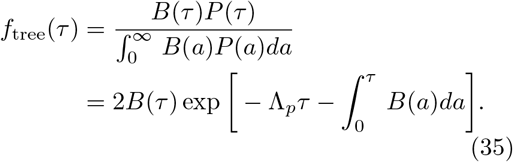

Reading now from the result for the lineage, Eq. (29), we obtain:

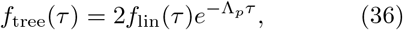

which corresponds to the result derived in Ref [10] with the identification of their generation time distribution *g* (resp. *g*^∗^) with our distributions *f*_lin_ (resp. *f*_tree_).

Using Eq. (36) we have, for instance:

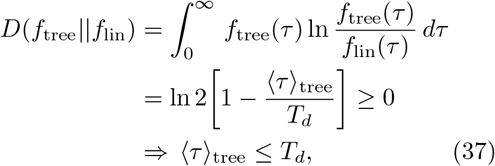

where as usual the population doubling time reads *T*_*d*_ = ln 2/Λ_*p*_. It is straightforward to prove the second inequality using the same technique. We then conclude that for IGT models, one has the same result as obtained for size structured populations in Eq. (25), i.e.,

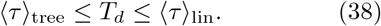

### D. Extension to age models with correlations

In view of the result of previous section, it is then natural to ask what happens in the more complex case in which mother-daughter correlations are present. In appendix B, we also derived a generalization of Eq. (36) for that case, namely:

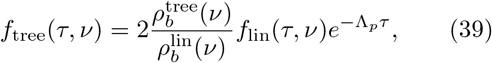

where 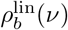 (resp. 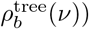) represent the growth rate distributions of newborn cells at the lineage (resp. tree level). The presence of these two new probability distributions is entirely due to mother-daughter correlations. As a result, the inequalities (25) (identical to (38)) do not necessarily hold for age models with correlations. An example where they are indeed violated can be found in the model with correlated generation times studied in Ref. [19] in some range of parameters.

## V. CONCLUSION

In conclusion, we have established several fluctuation relations which relate observables measured at the lineage and population levels. We have deduced from the second relation that mean generation times in a lineage should be larger than the population doubling times in models with cell size control, whether or not mother-daughter correlations are present, and in age models without correlations. In constrast to this, Eq. (25) can be violated in age models with correlations. Recent experiments reporting mean generation times in a lineage larger than the population doubling time [10], provide an illustration of the right inequality of Eq. (25). Our analysis indicates that such an observation is compatible with a cell size control model or with an age model without mother-daughter correlations. Further experimental tests of both inequalities of Eq. (25) based on our framework could reveal additional information on the underlying mechanism of cell size control.

Our approach being general, it could be extended to cover more complex cases such asymmetric divisions relevant for yeast cells, nonexponential regimes of growth, relevant for eukariots and other mechanisms of cell aging [25]. While we have mainly focused on the control of the size variable, extension of this formalism to other variables not directly linked to cell size is possible, one choice being for instance the protein copy numbers [20].

We also find that the variability of single cell growth has a negative impact on the population growth rate in the absence of mother-daughter correlations when the division rate is *B*(*x*, *ν*) = *νx*. A positive impact due to correlations has been reported in some other study [26], while more generally a positive or negative impact should be expected depending on the form of the division rate [23]. All these recent results suggest that generation times are under a strong evolutionary pressure in which single cell variability and correlations over generations [27] play an important role.

In the future, we would like to study systems where the division rate is controlled simultaneosly by the size and the age of the cell, which represents a situation of major biological relevance [28]. Finally, while this work was under review, two new studies of cell growth dynamics have appeared, which relate either to our pathwise formulation [29] or to our analysis of generation time distributions [30].

## ACKNOWLEDGMENTS

R.G.G. was supported by the Agence Nationale de la Recherche (ANR-16-CE11-0026-03) and by Labex CelTisPhysBio (ANR-10-LBX-0038). We would like to thank E. Braun, L. Robert, T. Nemoto, A. Olivier and T. J. Kobayashi for stimulating discussions.

## Appendix A: Path integral representation of the dynamics for size-controlled models

## 1. Population level

Let us start by building a path integral representation associated to the evolution of the number density of cells in the population case, Eq. (1). Here, we will allow for an arbitrary number of offsprings *m* for generality, although only *m* = 2 was considered above. We emphasize that *m* should be independent of the state of the system. Let us treat the following term

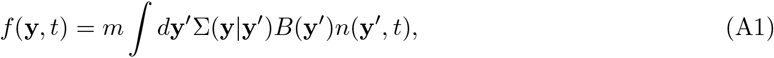

in Eq. (1) as a perturbation. The *growth propagator G*_*B*_ of the unperturbed dynamics is such that

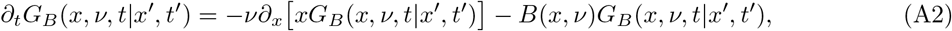

with initial condition *G*_*B*_ (*x*, *ν*, *t′*|*x′*, *t′*) = *δ*(*x − x′*). Then using these equations, one can check that

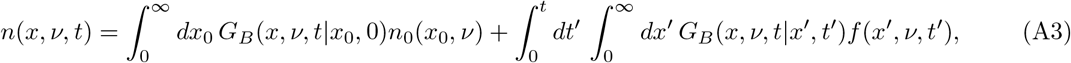

is equivalent to the initial problem given in Eq. (1). By explicitly using the definition of *f* from Eq. (A1), one obtains

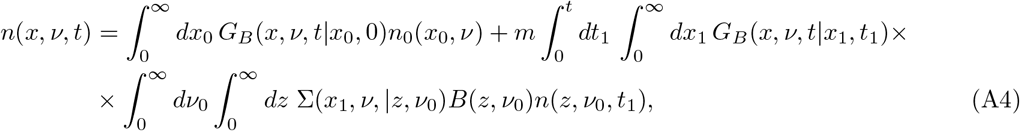

which allows to find an explicit solution for *n*(*x*, *ν*, *t*) iteratively.

The explicit solution of Eq. (A2) is

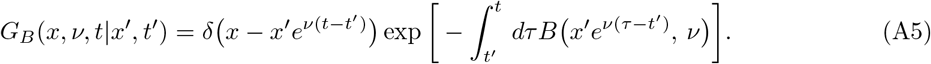

which allows us to write:

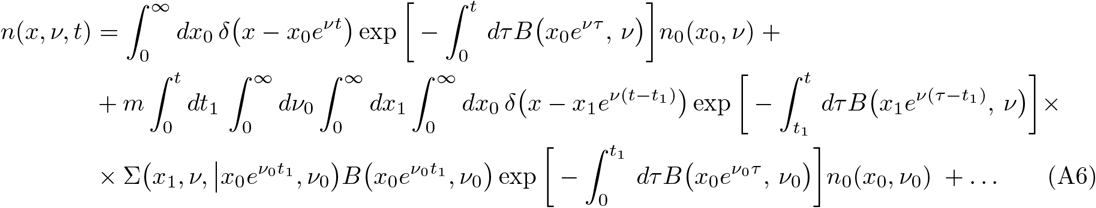

or more compactly:

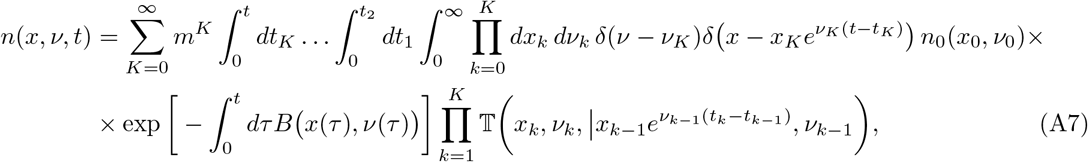

where trajectories explicity appearing in the exponential in the r.h.s. of (A7) are given as *ν*(*τ*) = *ν*_*k*_, and *x*(*τ*) = *x*_*k*_ exp(*ν*_*k*_(*τ − t*_*k*_)), for *τ* ∈ (*t*_*k*_, *t*_*k*+1_], while *k* = 0, 1, …, *K*. In our notations, *t*_0_ = 0 and *t*_*K*+1_ = *t*. In addition, the transition matrix is given as 𝕋(*x*, *ν*|*x′*, *ν′*) = Σ(*x*, *ν*|*x′*, *ν′*)*B*(*x′*, *ν′*).

The last step now consists in noticing that the object propagating trajectories from *t*_0_ = 0 up to time *t* in (A7) is not yet a path probability because it is not properly normalized. To deal with this issue it is good to pass from number densisites to population-level probability densities, *P* (*x*, *ν*, *t*) = *N* (*t*)^−1^*n*(*x*, *ν*, *t*), and *P*_0_(*x*, *ν*) = *N*(0)^−1^*n*_0_(*x*, *ν*). We can now write in terms of these quantities:

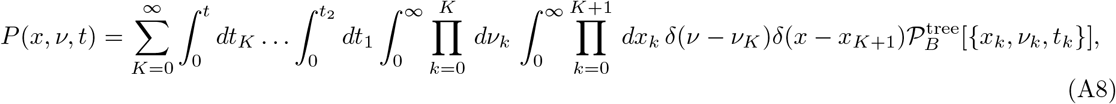

where the object

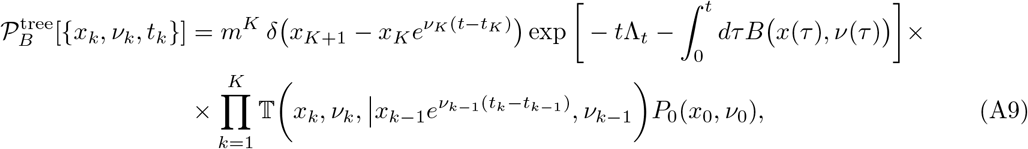

is now properly normalized and can be identified with the correct path propability generating averages of all observables related to the number density at the population level. We have added a subscript *B* to indicate that the division rate is given by *B*(*x*, *ν*). This will be important later in the derivation of fluctuation theorems. Note that when passing from densities to probability densities, a new term has appeared in the argument of the exponential namely Λ_*t*_, which is connected to the population growth rate, Λ_*p*_, by Eq. (11).

## 2. Lineage level

The starting point to derive the path probability for lineage observables is the evolution equation for the probability density of size and growth rate, Eq. (2) Except for the absence of the factor two in front of the integral, the structure of the equations are the same and the derivation follows along exactly as in the population case. We provide the final result:

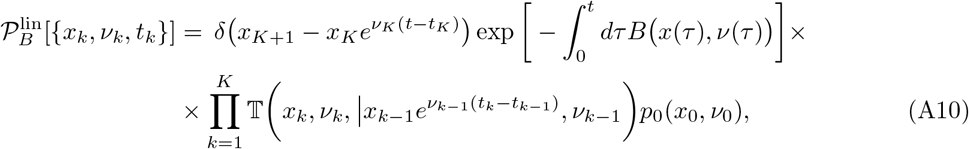

which can be readily shown to be properly normalized. Here *p*_0_ is the distribution of initial conditions for the lineage. Note that we have introduced *p*_0_, which could be different from the *P*_0_ introduced earlier as the initial condition of the population.

## 3. Derivation of fluctuation relations

We can now compare path probabilities representations at the population and lineage levels given by (A10) and (A9) with each other. We see that a possible way to bring both distributions “closer” together, is to multiply the division rate at the lineage level by the factor *m*, and to consider a lineage starting from the same initial condition as that of the population. A possible choice of this initial condition consists, for instance, in considering a population dynamics starting from a single cell.

In that case we have:

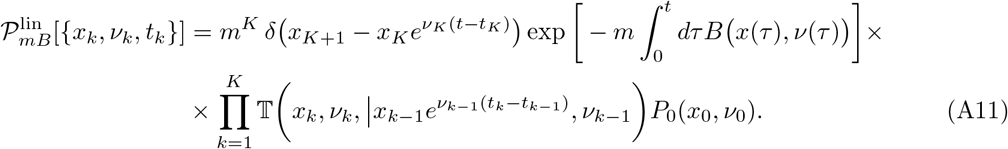

Then, the following relation holds from direct comparison of (A11) and (A9):

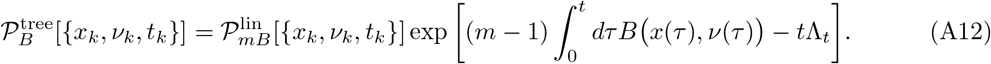

## Appendix B: Fluctuation theorem for correlated age models

## 1. Lineage dynamics

We now consider models in which interdivision times are correlated. The natural way in which these correlations arise is by inter-cell-cycle growth-rate fluctuations, as given by Eqs. (9) and (10). Growth-rate correlations are encoded in Σ, which is a properly normalized conditional probability. Again, we will focus on stationary conditions. It is simple to see from (9) that one can formally write the stationary distribution as:

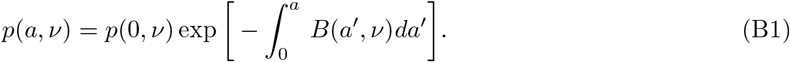

To determine *p*(0, *ν*), we use (10) and (B1):

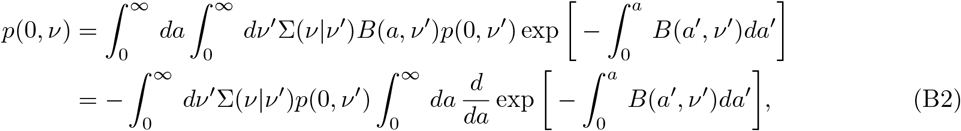

from where we get the following integral equation:

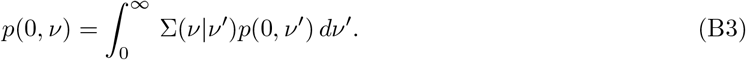

The generation time distribution can now be determined, again, as the age-distribution of dividing cells. It is worth considering slightly more general object, i.e., the *joint* probability distribution of interdivision time and growth rate

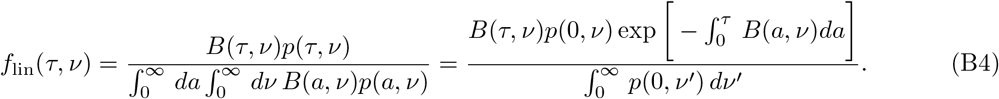

This result can be written in a more illuminating way by noticing that the growth rate distribution of newborn cells can be identified as

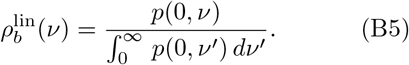

Furthermore, due to the linearity of Eq. (B3), and the fact that 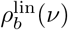 differs from *p*(0, *ν*) only in a multiplicative constant, we have that 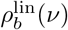 satisfies

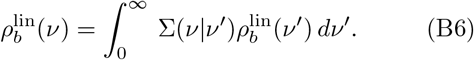

These observations then lead to the final result:

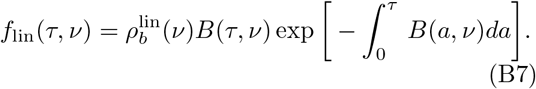

## 2. Population dynamics

Let us now consider the population level. The stationary equation for the population age distribution reads

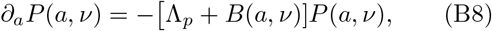

with boundary condition

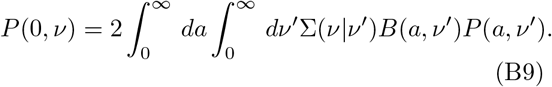

We then have

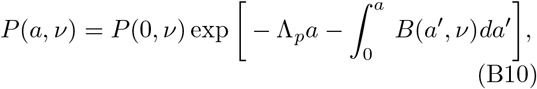

Note that the normalization of *P* gives the following condition:

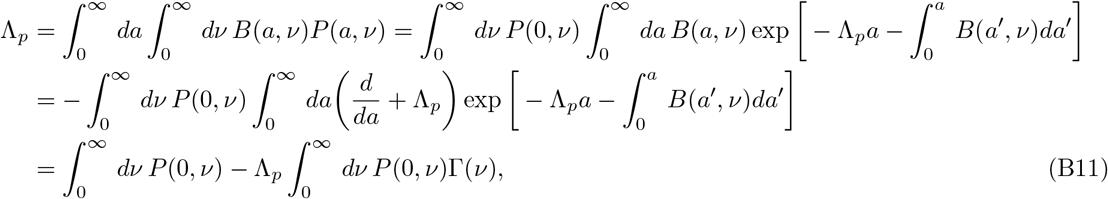

where we have used (B10) and introduced the function

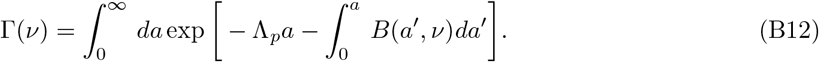

We can thus write for the growth rate of the population:

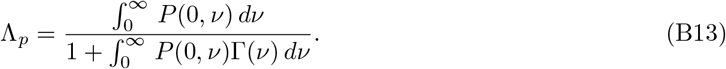

On the other hand, integrating directly in (B10), we get

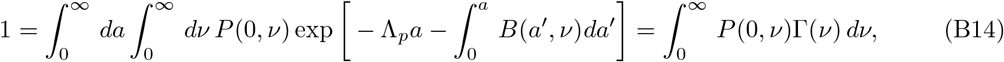

so we have

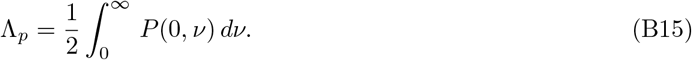

As before, we can find an equation for *P* (0, *ν*) using (B9) and the solution for *P* (*a*, *ν*), Eq. (B10):

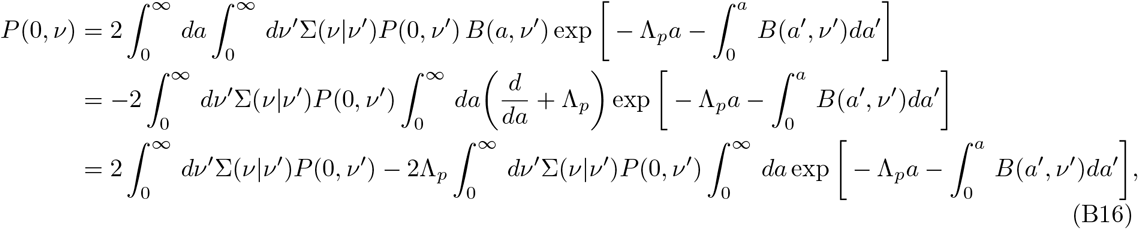

so, we then have

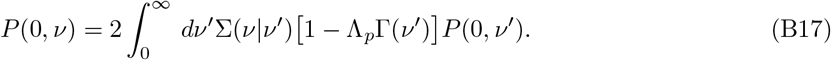

Let us now write the joint probability distribution of interdivision times and single-cell growth rate:

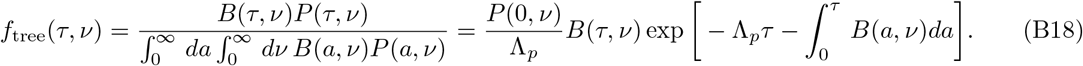

The condition (B15) implies that 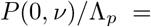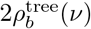, where

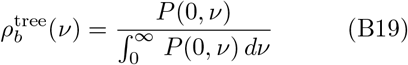

can be identified, as we did in the lineage case, with the growth rate distribution of newborn cells, now at the tree level. We then have:

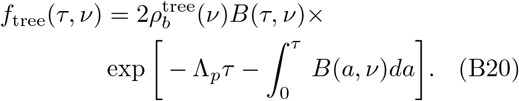

Note once more that the linearity of Eq. (B17) and the fact that *P* (0, *ν*) and 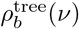 differ only on a multiplicative factor, lead to the equation satisfied by 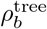:

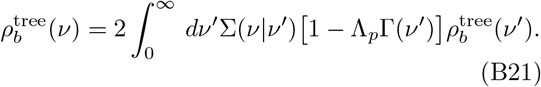

If we now compare (B20) and (B7), we readily obtain Eq. (39). Before closing this paragraph some comments are in order. First, note that as Eqs. (B6) and (B21) are clearly different, one has 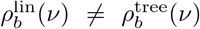. Nevertheless, in absence of fluctuations, when Σ(*ν*|*ν*′) = *δ*(*ν* − *ν*′), we have 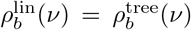. To illustrate this, let us consider, for instance, a population starting from a single cell with growth rate *ν*_0_. As the growth rate remains the same in all cell cycles, we have 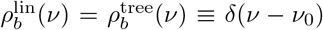 at all times. Then, Eq. (B6) becomes tautological, while Eq. (B21) leads to the identity

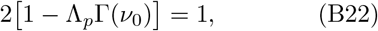

which is precisely the relation (34) found for IGT models (recall the definition of Γ, (B12)).

## 3. Inequalities in correlated age models

Let us now analyze the consequences of the generalized relation (39) for the inequalities. We have, for instance:

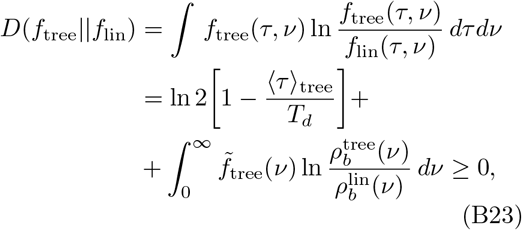

where 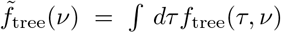 is the marginal distribution of the growth rate of the *dividing* cells. This result implies, in particular, that

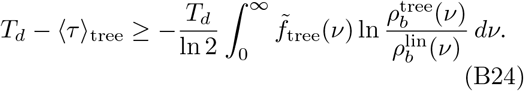

Given that the quantity in the right hand side of (B24) does not have a definite sign (in particular, it is not necessarily positive), in this case the left inequality in (38) (and Eq. (25)) may be violated. Repeating a similar argument, on arrives to the same conclusion for the right inequality.

## References

[1] A. Koch, Bacterial growth and form (Springer, 2001).

[2] L. Willis and K. Huang, Nat. Rev. Microbiol. 15, 606 (2017).

[3] A. Amir, Phys. Rev. Lett. 112, 208102 (2014).

[4] L. Robert, M. Hoffmann, N. Krell, S. Aymerich, J. Robert, and M. Doumic, BMC Biol. 12, 17 (2014).

[5] S. Taheri-Araghi, S. Bradde, J. T. Sauls, N. S. Hill, P. A. Levin, J. Paulsson, M. Vergassola, and S. Jun, Curr. Biol. 25, 385 (2015).

[6] J. Grilli, M. Osella, A. S. Kennard, and M. C. Lagomarsino, Phys. Rev. E 95, 032411 (2017).

[7] M. Campos, I. V. Surovtsev, S. Kato, A. Paintdakhi, B. Beltran, S. E. Ebmeier, and C. Jacobs-Wagner, Cell 159, 1433 (2014).

[8] E. O. Powell, Microbiology 15, 492 (1956).

[9] M. Kimmel and D. E. Axelrod, Branching Processes in Biology (Springer, 2015).

[10] M. Hashimoto, T. Nozoe, H. Nakaoka, R. Okura, S. Akiyoshi, K. Kaneko, E. Kussell, and Y. Wakamoto, Proc. Natl. Acad. Sci. 113, 3251 (2016).

[11] A. J. Hall, Steady Size Distributions in Cell Populations, Ph.D. thesis, Massey University (1991).

[12] T. J. Kobayashi and Y. Sughiyama, Phys. Rev. Lett. 115, 238102 (2015).

[13] Y. Sughiyama, T. J. Kobayashi, K. Tsumura, and K. Aihara, Phys. Rev. E 91, 032120 (2015).

[14] G. E. Crooks, Phys. Rev. E 61, 2361 (2000).

[15] T. Nemoto, F. Bouchet, R. L. Jack, and V. Lecomte, Phys. Rev. E 93, 062123 (2016).

[16] C. Giardinà, J. Kurchan, and L. Peliti, Phys. Rev. Lett. 96, 120603 (2006).

[17] V. Bansaye, J.-F. Delmas, L. Marsalle, and V. C. Tran, Ann. Appl. Probab. 21, 2263 (2011).

[18] T. Nozoe, E. Kussell, and Y. Wakamoto, PLoS. Genet. 13, 1 (2017).

[19] J. Lin and A. Amir, Cell Systems 5, 358 (2017).

[20] N. Brenner, E. Braun, A. Yoney, L. Susman, J. Rotella, and H. Salman, Eur. Phys. J. E 38, 102 (2015).

[21] M. R. Evans and S. N. Majumdar, Phys. Rev. Lett. 106, 160601 (2011).

[22] M. Doumic, B. Perthame, and J. P. Zubelli, Inverse Probl. 25, 045008 (2009).

[23] A. Olivier, Am. Inst. of Math. Sci. 10, 481 (2017).

[24] Y. Wakamoto, A. Y. Grosberg, and E. Kussell, Evolution 66, 115 (2011).

[25] E. J. Stewart, R. Madden, G. Paul, and F. Taddei, PLOS Biology 3 (2005).

[26] A. Amir and J. Lin, “Population growth with correlated generation times at the single-cell level,” (2018), arxiv:1806.02818.

[27] O. Rivoire and S. Leibler, Proc. Natl. Acad. Sci. U.S.A. 111, 1940 (2014).

[28] M. Osella, E. Nugent, and M. Cosentino Lagomarsino, Proc. Natl. Acad. Sci. U.S.A. 111, 3431 (2014).

[29] N. S. Sughiyama, Y. and T. J. Kobayashi, ArXiv:1806.0021 (2018).

[30] F. Jafarpour, C. S. Wright, H. Gudjonson, J. Riebling, E. Dawson, K. Lo, A. Fiebig, S. Crosson, A. R. Dinner, and S. Iyer-Biswas, Phys. Rev. X 8, 021007 (2018).

